# *FADS1* and the timing of human adaptation to agriculture

**DOI:** 10.1101/337998

**Authors:** Sara Mathieson, Iain Mathieson

## Abstract

Variation at the *FADS1/FADS2* gene cluster is functionally associated with differences in lipid metabolism and is often hypothesized to reflect adaptation to an agricultural diet. Here, we test the evidence for this relationship using both modern and ancient DNA data. We show that almost all the inhabitants of Europe carried the ancestral allele until the derived allele was introduced approximately 8,500 years ago by Early Neolithic farming populations. However, we also show that it was not under strong selection in these populations. We find that this allele, and other proposed agricultural adaptations at *LCT/MCM6* and *SLC22A4*, were not strongly selected until much later, perhaps as late as the Bronze Age. Similarly, increased copy number variation at the salivary amylase gene *AMY1* is not linked to the development of agriculture although, in this case, the putative adaptation precedes the agricultural transition. Our analysis shows that selection at the *FADS* locus was not tightly linked to the initial introduction of agriculture and the Neolithic transition. Further, it suggests that the strongest signals of recent human adaptation in Europe did not coincide with the Neolithic transition but with more recent changes in environment, diet or efficiency of selection due to increases in effective population size.

## Introduction

Human history has seen a number of major transitions in diet (Luca, et al. 2010). The most recent was the transition to a modern “industrialized” diet based on intensive farming and highly processed food. Before that, many parts of the world saw a dramatic transition from a diet based on hunting and gathering to a diet heavily based on the products of agriculture. The transition occurred independently several times in different parts of the world, but the earliest known example is the Fertile Crescent, at least 10,500 years ago. From there agriculture spread gradually northwest through Anatolia into Europe, and eastwards to the Indian subcontinent (Bellwood 2004). Even outside these periods of transition, differences in diet based on both cultural preferences and food source availability would have been a major aspect of environmental differences between human populations–differences that would likely lead to genetic adaptation. Thus, by identifying and studying the evolution of genetic adaptations to diet, we learn not only about historical changes in diet, but also about the genetic basis of diet-related phenotypic differences among present-day human populations.

One previously identified adaptation involves the fatty acid desaturase genes *FADS1* and *FADS2*. These genes encode proteins which catalyze key steps in the ω-3 and ω-6 lipid biosynthesis pathways (Nakamura and Nara 2004). These pathways synthesize long-chain (LC) polyunsaturated fatty acids (PUFA) necessary for cell- and, particularly, neuronal-membrane development from short-chain (SC) PUFA (Darios and Davletov 2006). The evolutionary interaction with diet stems from the fact that different diets contain different ratios of SC- and LC-PUFA. Specifically, diets that are high in meat or marine products contain relatively high LC-PUFA levels, and thus may require lower levels of *FADS1* and *FADS2* activity compared to diets that are high in plant-based fats (Ameur, et al. 2012; Mathias, et al. 2012; Fumagalli, et al. 2015; Kothapalli, et al. 2016; Buckley, et al. 2017; Ye, et al. 2017). As a result of this environmental interaction, these genes have been repeatedly targeted by natural selection.

Most dramatically, a derived haplotype (“haplotype D”, following Ameur, et al. (2012)) containing the 3’ end of *FADS1* is at very high frequency in present-day African populations and intermediate to high frequency in present-day Eurasians (Ameur, et al. 2012; Mathias, et al. 2012), and experienced ancient positive selection in Africa (Ameur, et al. 2012; Mathias, et al. 2012). The derived haplotype increases expression of *FADS1* (Ameur, et al. 2012) and likely represents an adaptation to a high ratio of SC- to LC-PUFA, *i.e*. to a plant-based diet. In Europe, direct evidence from ancient DNA has shown that haplotype D was rare around 10,000 years before present (BP) but has increased in frequency since then, likely due to selection, to its present-day frequency of ~60% in Europe (Mathieson, et al. 2015). This increase was plausibly associated with the adoption – starting around 8,500 BP in Southeastern Europe before spreading North and West – of an agricultural lifestyle and diet that would have a higher SC- to LC-PUFA ratio than the earlier hunter-gatherer diet (Mathieson, et al. 2015; Buckley, et al. 2017; Ye, et al. 2017). Interestingly, the Altai Neanderthal genome shares at least a partial version of haplotype D (Buckley, et al. 2017; Harris, et al. 2017), suggesting that the functional variation at this locus may predate the split of Neanderthals and modern humans.

Other haplotypes at the locus have been shown to be under selection in different populations in more recent history. In particular, another haplotype that is common in, but largely restricted to, the Greenlandic Inuit population reduces the activity of *FADS1*, is associated with PUFA levels, and is likely an adaptation to a diet that is extremely high in LC-PUFA from marine sources (Fumagalli, et al. 2015). Conversely, a variant that increases expression of *FADS2* has been selected in South Asian populations – and may be a specific adaptation to a vegetarian diet (Kothapalli, et al. 2016). Reflecting its important role in lipid metabolism, variation at the *FADS* locus also contributes significantly to variation in lipid levels in present-day populations. As well as directly contributing to variation in PUFA levels, SNPs in haplotype D are among the strongest genome-wide association study (GWAS) signals for triglyceride and cholesterol levels (Teslovich, et al. 2010). Thus, the complex evolutionary history of the region is not only informative about ancient human diets, but also potentially relevant for understanding the distribution of lipid-related disease risk both within and between populations. We therefore aimed to characterize the history of the region, by combining inference from ancient and modern DNA data, in order to understand the evolutionary basis of this important functional variation and its relationship with changes in diet. We also compared the evolutionary history of the *FADS* locus with the histories of other loci involved in dietary adaptation, to see whether we could detect shared patterns of adaptation.

## Results

### Haplotype structure at *FADS1*

We began by investigating 300 high-coverage whole-genome sequences from the Simons Genome Diversity Project (SGDP) (Mallick, et al. 2016), as well as three archaic human genomes (Meyer, et al. 2012; Prufer, et al. 2014; Prufer, et al. 2017), to investigate the fine-scale haplotype structure in the region (Figure 1, Supplementary Figure 1). By clustering haplotypes using a graph-adjacency algorithm (Methods), we defined three nested “derived” haplotypes (Figure 1A, Table 1, Supplementary Table 1, Supplementary Figure 1). Haplotype D extends over 78kb, is largely restricted to Eurasia, and is comparable to the haplotype D defined by Ameur, et al. (2012). Haplotype C is an 18kb portion of haplotype D that is shared between African and Eurasian populations. Finally, haplotype B is a 15kb region that represents the portion of haplotype D that is shared between modern humans and both the Altai (Prufer, et al. 2014) and Vindija (Prufer, et al. 2017) Neanderthals. We denote the modern human haplotype carrying ancestral alleles at haplotype-defining SNPs as haplotype A (the “ancestral” haplotype). This analysis also allows us to prioritize likely causal SNPs, which are currently unknown (Buckley, et al. 2017; Ye, et al. 2017). If the derived SNP that was selected in modern humans was also selected in Neanderthals, then it must lie in haplotype B, which is defined by just four SNPs (rs174546, rs174547, rs174554 and rs174562).

**Figure 1:**
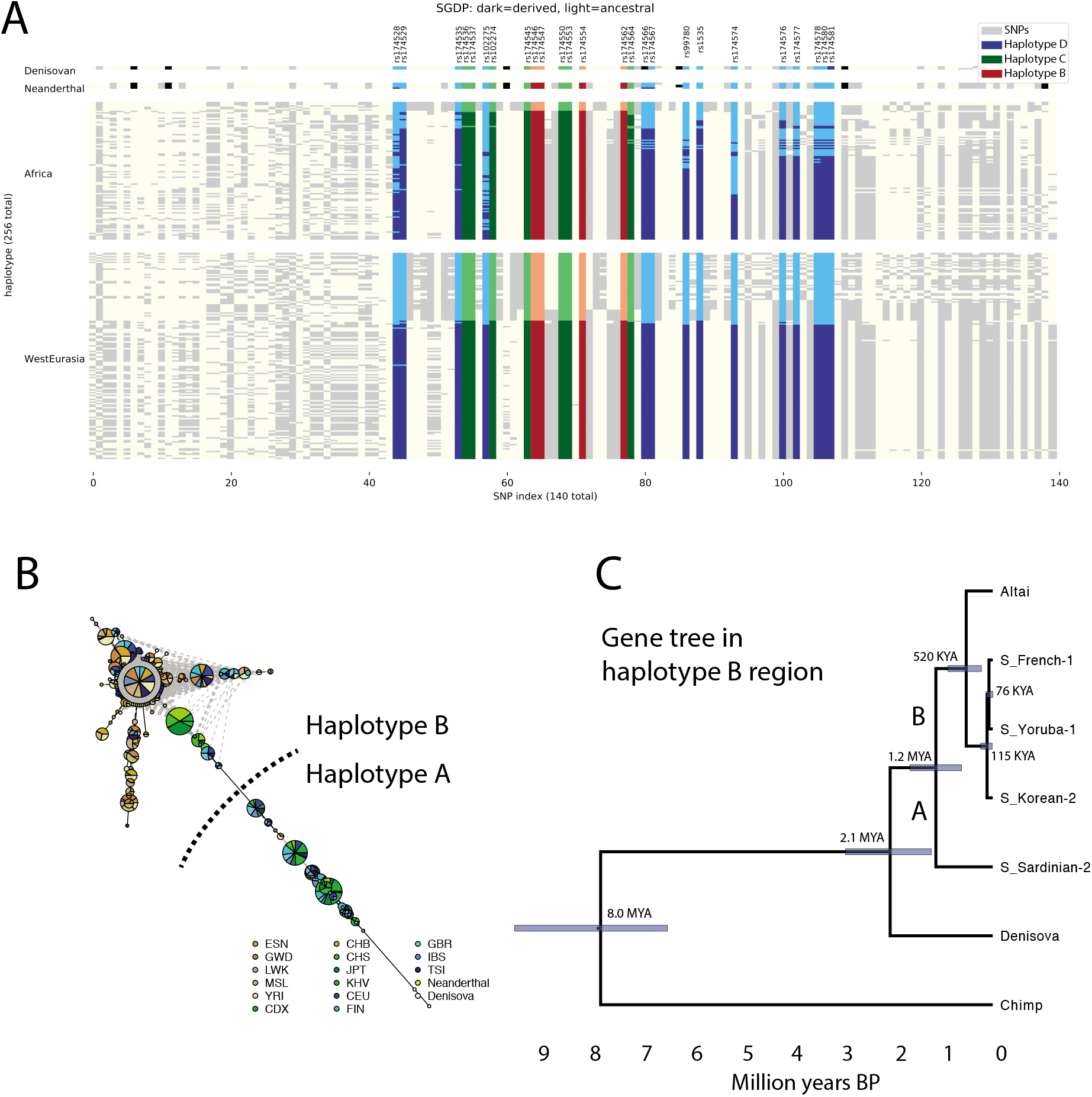
Haplotype structure at the *FADS1* region. **A**: Haplotypes from the SGDP African and West Eurasian populations (Mallick, et al. 2016) and the Neanderthal (Prufer, et al. 2014; Prufer, et al. 2017) and Denisovan (Meyer, et al. 2012) genomes. Each column represents a SNP and each row a phased haplotype. Dark and light colors represent derived and ancestral alleles at each SNP and blue, green and red colors indicate SNPs that are part of haplotypes D, C and B, respectively. **B**: Haplotype network for region B constructed from 1000 Genomes (1000 Genomes Project Consortium 2015) and archaic samples. Green, blue and brown indicate East Asian, European and African populations respectively. **C**: Gene tree for the haplotype B region inferred for representative haplotypes.

**Table 1:** Haplotypes described in the text. Summary of the 4 haplotypes discussed. See Supplementary Table 1 for defining SNPs.

| Haplotype | Length (kb) | State | Shared |
| --- | --- | --- | --- |
| **A** | - | Ancestral | - |
| **B** | 15 | Derived | Neanderthal and present-day humans |
| **C** | 18 | Derived | Present-day humans |
| **D** | 78 | Derived | Present-day Eurasians |

**Table 2:** Summary of ABC results for three possible episodes of selection. We inferred the probability of selection on a new mutation (as opposed to standing variation), estimated selection-onset time, 95% credible intervals for this time, and 95% credible intervals for the selection coefficient. See Supplementary Tables 2 and 3 for more detailed results.

|  | P(New mut.) | Start (yrs BP) | Start CrI (yrs BP) | Sel. Coeff. |
| --- | --- | --- | --- | --- |
| **1. Selection for the derived allele in the ancestors of present-day Africans** | 0.74 | 310,548 | 202,409-492,231 | 0.001-0.002 |
| **2. Selection for the ancestral allele in the ancestors of present-day Europeans** | 0.25 | 246,399 | 49,358-1,137,909 | 0.000-0.028 |
| **3. Selection for the derived allele in Europe** | 0.00 | 2,538 | 1,695-4,048 | 0.034-0.069 |

Using data from the 1000 Genomes project (1000 Genomes Project Consortium 2015), we replicate the observation (Mathias, et al. 2012; Buckley, et al. 2017; Harris, et al. 2017) that haplotypes in the haplotype B region fall into two clusters (Figure 1B, Supplementary Figure 2). One cluster contains Eurasian-ancestry individuals that carry haplotype A (and all individuals with Native American ancestry). The other cluster contains Neanderthals, Eurasians that carry haplotype D, and almost all present-day African-ancestry individuals. A small number of intermediate haplotypes are either recombinant or the result of phasing errors. We inferred the phylogeny of the haplotype B region using BEAST2 (Figure 1C) (Bouckaert, et al. 2014). The region is small and has a very low recombination rate, (average 0.03 cM/mb according to the International HapMap Consortium (2007) combined recombination map), so we ignored possible recombination events. Assuming that the common ancestor of human and chimp haplotypes was 6.6-10.0 million years ago – corresponding to a genome-wide mutation rate of ~4-6× 10^−10^ per-base per-year (Amster and Sella 2016; Scally 2016) – we infer that the most recent common ancestor (MRCA) of the present-day African (C) and European (D) haplotypes was around 76,000 BP, of haplotypes B and C around 520,000 BP, and that of haplotypes A and B around 1.2 million BP. These dates are consistent with the estimates of Harris, et al. (2017) and suggest that the European-specific haplotype D diverged around the time of the out-of-Africa bottleneck and the diversification of non-African lineages (Mallick, et al. 2016). They also suggest that modern human ancestral and derived haplotypes coexisted in the Denisovan-Neanderthal-modern human ancestral population, with the Neanderthal haplotype splitting off around the time of mean human-Neanderthal divergence 550-765,000 BP (Prufer, et al. 2014).

## ABC analysis of ancient selection in Africa and recent selection in Eurasia

Having established the structure and phylogeny of the haplotypes in the region, we used approximate Bayesian computation (ABC) (Wegmann, et al. 2010; Peter, et al. 2012) to infer the strength and timing of selection on the derived haplotype in Africa (Table 1, Supplementary Table 2), first described by Mathias, et al. (2012). Using data from the 1000 Genomes project, treating the derived allele of rs174546 as the selected SNP, and fixing the mutation rate to 1.25×10^−8^ per-base per-generation, we infer that selection most likely but not definitively acted on a new mutation (posterior probability of new mutation 0.74, range 0.13-0.99, see Peter, et al. (2012) for details of the inference approach) and estimate that selection began 202,000-492,000 BP (Supplementary Table 2). This estimates is uncertain though, with 95% credible intervals (CrIs) in different populations ranging from 84,000-1,426,000 BP. Mathias, et al. (2012) estimated 85,000 ± 84,000 years for the onset of selection in Africa. However, that analysis assumed a Human-Chimpanzee split of 6.5 million years, now thought to be an underestimate and, if appropriately rescaled, would overlap the low end of our estimate. Overall, the evidence suggests that selection began around or before the time of the deepest splits among human populations (Mallick, et al. 2016; Schlebusch, et al. 2017) 200,000-250,000 BP. This is consistent with the observation that the derived allele is shared among all present-day African populations including those, such as Khoe-San and Mbuti, which were substantially diverged from the ancestors of other present-day African populations at least 100,000 BP (Mallick, et al. 2016; Schlebusch, et al. 2017).

In contrast, when we analyze present-day European genomes, we find that strong (selection coefficient s=3.4-6.9%) selection for the derived allele acted on standing variation (posterior probability=1.0) much more recently – starting between 1700 and 4000 BP (95% CrIs in different populations range from 1000-12,000 BP, Supplementary Table 2). This recent selection is hard to reconcile with the suggestion (Mathieson, et al. 2015; Ye, et al. 2017) that it is closely linked to the development of agriculture, at least 10,500 BP (Bellwood 2004). It is also unclear why the derived haplotype was at such low frequency in pre-agricultural Europeans (Mathieson, et al. 2015; Ye, et al. 2017), when ABC results and derived allele sharing between African populations would imply that the derived allele would have been at high frequency before the split of African and non-African ancestral populations. To resolve these questions, we turned to ancient DNA data.

## Low frequency of the derived *FADS1* allele in Upper Palaeolithic Eurasia

We first investigated why the derived haplotype was at such low frequency in pre-agricultural Europeans, by analyzing data from 52 Paleolithic and Mesolithic individuals dated between 45,000 and 8,000 BP (Fu, et al. 2014; Jones, et al. 2015; Fu, et al. 2016; Sikora, et al. 2017). We infer the presence of haplotype D based on 5 SNPs that were typed on the “1240k” capture array used for most of these samples (Fu, et al. 2016). The derived haplotype is rare in all populations (Figure 2A, “Direct observations”). When we tried to use imputation to increase sample size, the results were inconsistent. Specifically, imputed data suggested a much higher frequency in most of the ancient population groupings, for example around 40% frequency in the Vestonice population. This higher frequency agrees with a previous analysis of this data by Ye, et al. (2017). However, we find that imputation is unreliable for these data because many of the Ice Age samples have extremely low coverage. This leads to a high rate of false positive inference of the derived allele because it is at relatively high frequency in the present-day populations used as a reference panel. Older samples tend to have lower coverage and thus higher false positive rates, leading to a spurious inference of a decline in frequency over time. We estimated the false positive rate as a function of coverage by simulating low coverage data from present-day samples that carry the ancestral allele. Non-zero derived allele frequency estimates in the imputed data are consistent with the estimated false positive rate (Methods, Figure 2B). In fact, we find reliable evidence of the derived allele in only one individual – from the 34,000 BP site of Sunghir (Sikora, et al. 2017). Therefore, it seems likely that the derived allele was rare among early “out-of-Africa” populations, at least in Western Eurasia.

**Figure 2:**
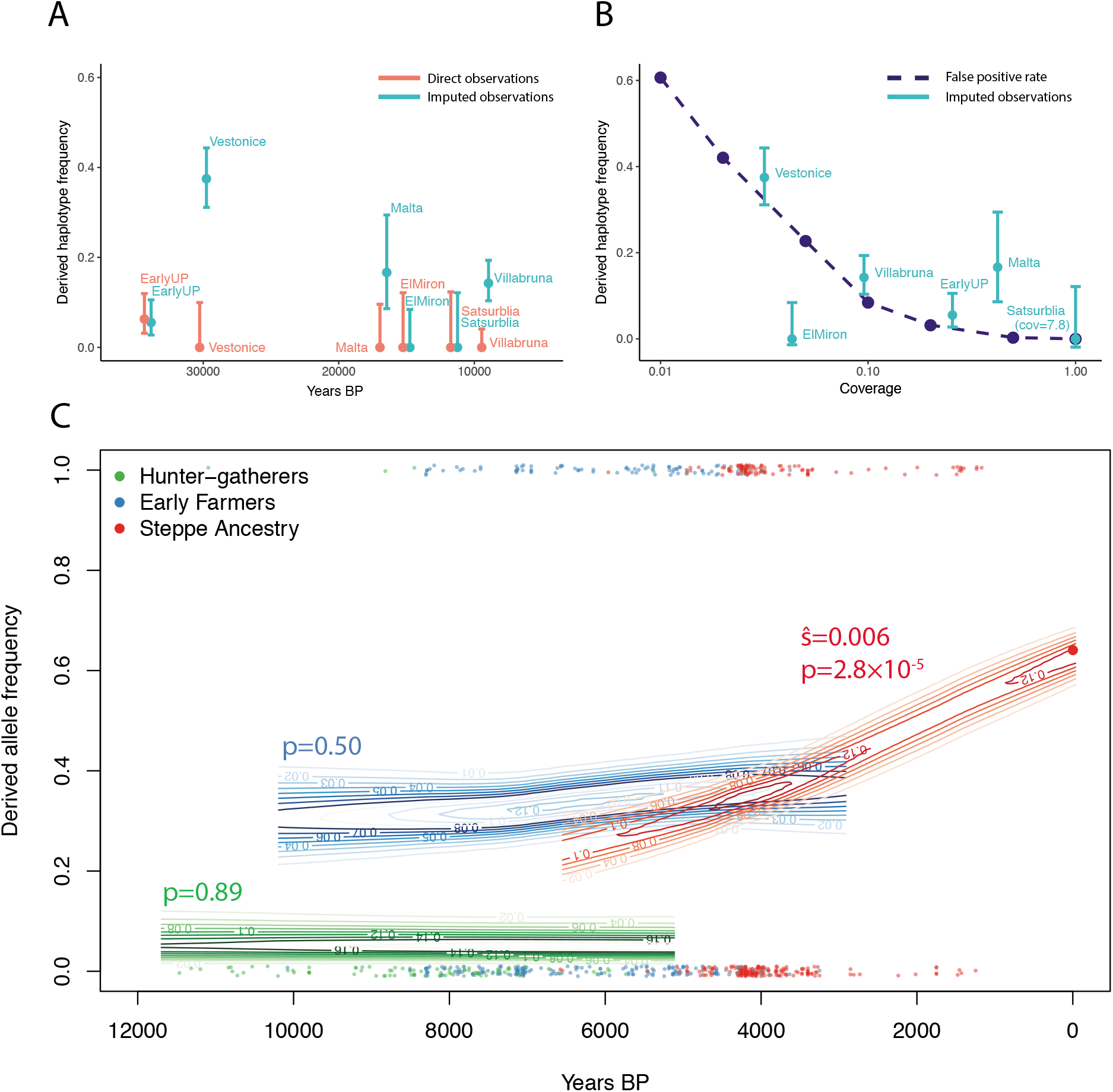
Direct ancient DNA evidence for the history of *FADS1*. **A**: Derived haplotype frequency estimated from direct observation of SNPs on the haplotype (red) and imputed data (blue) in Upper Palaeolithic individuals (45,000-10,000 BP) (Fu, et al. 2016). **B**: Estimated imputation false positive rate as a function of coverage (dashed line). Imputed allele frequencies in Upper Palaeolithic populations plotted for comparison at the median coverage in that population. **C**: Allele frequencies (at rs174546) over the past 12,000 years estimated from 1055 (669 with coverage) ancient and 99 modern individuals. Each point is an ancient pseudo-haploid individual call, at the bottom of the plot if it is ancestral and the top if it derived. Contours indicate the posterior probability of allele frequencies in the ancient populations and P-values for non-zero selection coefficients are indicated.

This lack of the derived allele in early non-Africans is surprising because our ABC analysis suggested that selection within the ancestral population began at least 200,000 years ago, and the derived allele would therefore be expected to have been at high frequency when African and non-African ancestors diverged. Further, the derived allele is shared between present-day African populations that were genetically isolated before the split of present-day African and non-African ancestors (Mallick, et al. 2016; Schlebusch, et al. 2017). Ye, et al. (2017) determined that the ancestral allele must have been selected in the ancestors of present-day Europeans, but located this selection in Europe, after the out of Africa bottleneck. Our analysis suggests that in fact, this selection was much older, occurring either before or during the bottleneck. To test this, we used the same ABC approach we used to investigate the derived allele to infer selection on the ancestral allele (Methods, Table 1, Supplementary Table 2). We infer that the ancestral allele was selected from standing variation in the ancestors of present-day Europeans (s=0.1%, 95% CrI=0.0-2.8%), starting around 246,000 BP (95% CrI 49,000-1,138,000 BP). This large credible interval overlaps the credible intervals for selection on the derived allele in African ancestors and means that we do not have power to resolve these selective events. We also estimated the timing of selection using an independent approach based on haplotype decay (Smith, et al. 2017), qualitatively supporting ancient selection for the derived allele (Supplementary Table 4). These results support the observation that if there was selection on the ancestral allele in the ancestors of present-day Europeans, it occurred mostly before the date of the earliest ancient samples with genetic data.

## Selection for the derived *FADS1* allele in Europe was not closely linked to early agriculture

The derived haplotype was almost absent in Upper Palaeolithic and Mesolithic Europe (before ~8,500 BP), but is today at a frequency of ~60%. Previous analysis of both ancient and modern DNA has inferred strong selection for the allele over the past 10,000 years (Mathieson, et al. 2015; Field, et al. 2016; Buckley, et al. 2017; Ye, et al. 2017). This has been interpreted to mean that selection for the derived allele was driven by the development of agriculture, around 10,500 BP–a reasonable interpretation since the derived allele is plausibly an adaption to a diet high in plant fats and low in animal fats. However, our ABC analysis suggests that selection in Europe might actually be restricted to the past few thousand years and thus post-date the development of agriculture by many millennia. To test this directly, we analyzed data from 1055 ancient Europeans who lived between 12,000 and 1,000 BP (Keller, et al. 2012; Fu, et al. 2014; Gamba, et al. 2014; Lazaridis, et al. 2014; Olalde, et al. 2014; Raghavan, et al. 2014; Seguin-Orlando, et al. 2014; Allentoft, et al. 2015; Gunther, et al. 2015; Haak, et al. 2015; Jones, et al. 2015; Mathieson, et al. 2015; Cassidy, et al. 2016; Hofmanová, et al. 2016; Martiniano, et al. 2016; Schiffels, et al. 2016; Jones, et al. 2017; Lazaridis, et al. 2017; Lipson, et al. 2017; Mathieson, et al. 2018; Olalde, et al. 2018). We divided these individuals into three populations, based on their genome-wide ancestry (Methods). First, individuals with “hunter-gatherer ancestry” were the Mesolithic inhabitants of Europe or their descendants. Second, “Early Farmers” were people from Neolithic Northwest Anatolia or their descendants, possibly admixed with hunter-gatherers, who migrated throughout Europe. Finally, people with “Steppe ancestry” had ancestry that was originally derived from Bronze Age steppe populations like the Yamnaya. (Lazaridis, et al. 2014; Allentoft, et al. 2015; Haak, et al. 2015). Because transitions between these populations involved dramatic genetic discontinuity, with 75-100% of ancestry replaced (Allentoft, et al. 2015; Haak, et al. 2015), we analyzed each of them separately.

In each of these three populations, we estimated the frequency and selection coefficient of the derived allele by fitting a hidden Markov Model (HMM) to the time series of observations (Methods, Figure 2C) (Mathieson and McVean 2013). We estimate that the selection coefficient in both hunter-gatherer and Early Farmer populations was not significantly different from zero (i.e. no evidence that the allele was under selection), and a selection coefficient of 0.6% (95% CI 0.4-1.5%) in Steppe-ancestry populations. This selection coefficient is lower than that estimated using ABC. One possible explanation is that the HMM forces the selection coefficient to be constant in each population, so if selection operated for only part of the time represented, or in a subset of the population, its strength might be underestimated. However, both the ancient DNA and ABC analyses are consistent with a relatively recent onset of selection and show that, although the derived allele was present in early farming populations, it was not strongly selected. Plausibly, the derived allele was introduced to the ancestors of the early farmers through admixture with a population that carried “basal Eurasian” ancestry (Lazaridis, et al. 2016) not found in Palaeolithic or Mesolithic Europe. Alternatively, the allele may have been retained in some, unsampled, Upper Paleolithic Eurasian population, or driven to higher frequency somewhere by one or more additional ancient episodes of selection.

Because agriculture was introduced to different parts of Europe at different times, spreading broadly from south to north, we split the ancient DNA data into Northern and Southern European groups (Methods) and re-ran the analysis, obtaining similar results in both groups (Supplementary Figure 3). To confirm the key observation that the derived allele was not under selection in Early Farmers, we restricted to this group and fitted a logistic regression to the observations with date, hunter-gatherer ancestry, Steppe ancestry and latitude as covariates. None of these four coefficients were significantly different from zero (P=0.19, 0.89, 0.41 and 0.78 respectively), confirming that selection for the allele is not being masked by variation in ancestry (for example by increasing hunter-gatherer ancestry over time), or in the geographic position of sampled individuals (for example because agriculture began at different dates at different latitudes). Finally, we split the Early Farmer and Steppe ancestry populations into 4000 year chunks and analyzed them separately (Supplementary Figure 4). The frequency of the derived allele is higher in later Early Farmers (after 6,000 BP), compared to early Early Farmers (40% vs 29%, Fisher’s exact test P=0.026), perhaps reflecting a shift in ancestry. But we find no evidence of selection in either early or later Early Farmers, or in early Steppe ancestry populations (before 4,000 BP).

## Patterns of population differentiation at other lipid-associated alleles

SNPs that tag the derived haplotype are among the strongest genome-wide associations with lipid levels (Teslovich, et al. 2010). To test whether the selective events we observe were consequences of more general selection on lipid levels, we investigated patterns of African-European population differentiation between variants associated with three blood lipid traits–triglycerides (TG), high-density cholesterol (HDL) and low-density cholesterol (LDL) (Teslovich, et al. 2010). We find that, for HDL and TG, trait-increasing alleles tend to be more common in African than European populations, while for LDL, the trait increasing allele is more common in European populations (Figure 3A, Methods). These effects are in the opposite direction to those of the *FADS1* haplotype. The derived allele, which is more common in African than European populations, tends to decrease HDL and TG and increase LDL. We conclude that selection on *FADS1* was not driven by its effect on overall blood lipid levels (which, if anything, moved in the opposite direction), but by its specific effect on PUFA synthesis. We further find that there is no significant difference in the frequency of LDL-increasing alleles when comparing the three ancient populations (Figure 3B). Since they do differ in the frequency of the *FADS1* allele, this suggests that recent selection on *FADS1* was also not driven by selection more generally on lipid levels.

**Figure 3:**
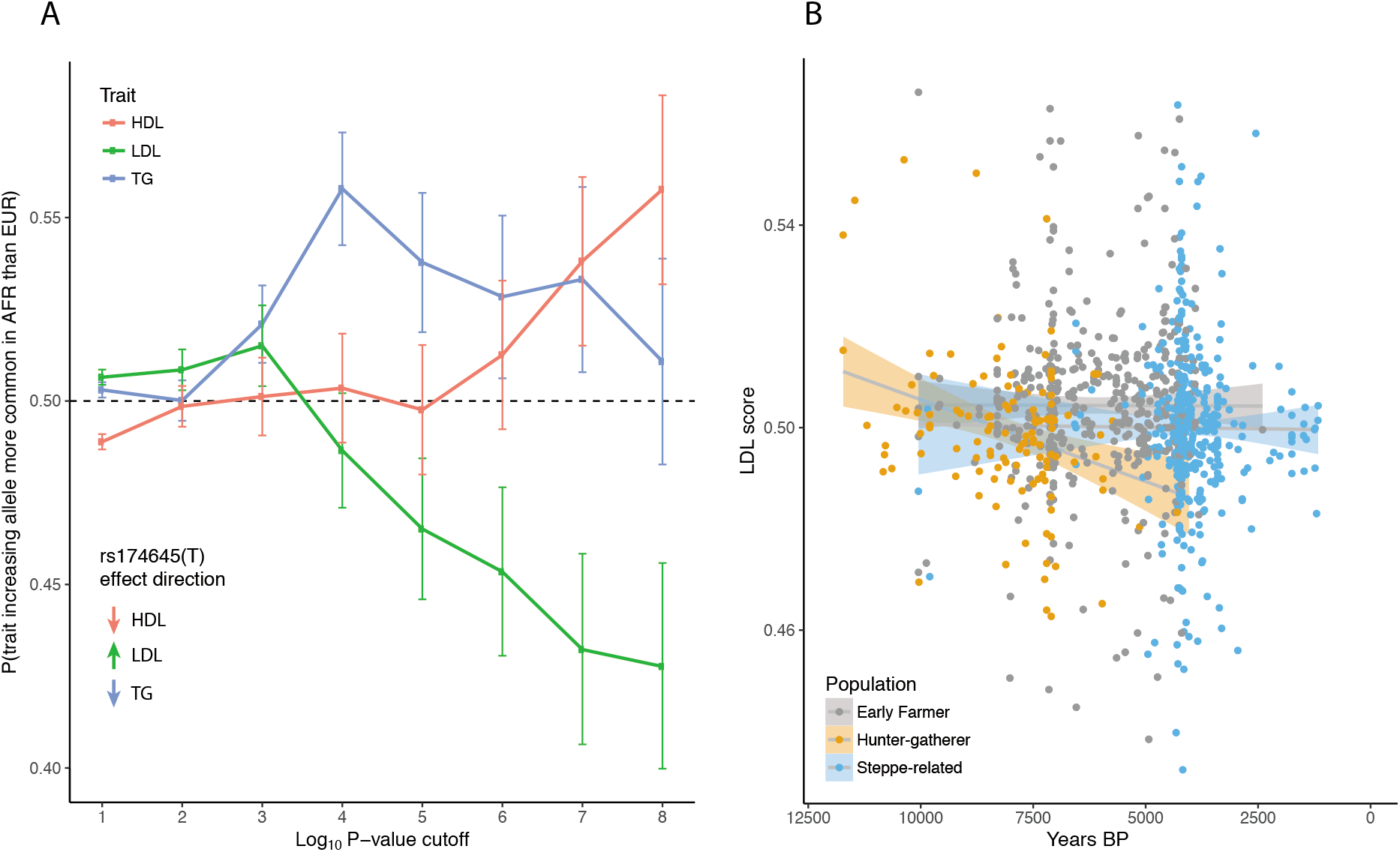
Trends in lipid-associated allele frequencies. **A**: Probability that the trait-increasing allele is more common in African than European populations in the 1000 Genomes Project data (1000 Genomes Project Consortium 2015) for lipid-related traits. We show results for each trait with different P-value cutoffs. Vertical bars represent 95% confidence intervals. Inset shows the direction of effect for the derived FADS1 haplotype, using rs174546 as the tag SNP. **B**: LDL score for 1003 ancient individuals, classified according to ancestry, as a function of their age. LDL score is the proportion of significant LDL-associated variants at which the ancient individual carries the trait-increasing allele (Methods).

## Patterns of population differentiation at other diet-associated variants

We investigated whether other variants that have been suggested to be associated with the adoption of agriculture showed similar temporal patterns of selection to *FADS1*. The salivary amylase gene *AMY1* is highly copy-number variable among present-day populations, ranging from a diploid copy number of 2 (the ancestral state) to 17 (Groot, et al. 1991; Perry, et al. 2007; Usher, et al. 2015), with a mean of 6.7 copies in present-day Europeans (Usher, et al. 2015). It has been suggested that increased copy number improves the digestion of starchy food and is therefore an adaptation to an agricultural diet that is relatively rich in starch (Perry, et al. 2007). To date, one Early Farmer has been reported to have a relatively high *AMY1* copy number of 16 (Lazaridis, et al. 2014), while six hunter-gatherers had 6-12 copies (Lazaridis, et al. 2014; Olalde, et al. 2014; Gunther, et al. 2018). Inchley, et al. (2016) showed that the initial expansion of *AMY1* copy number predated the out-of-Africa bottleneck, and found no evidence of recent selection. To investigate this directly, we called *AMY1* copy number in 76 ancient West Eurasian individuals with published shotgun sequence data (Keller, et al. 2012; Meyer, et al. 2012; Skoglund, et al. 2012; Gamba, et al. 2014; Lazaridis, et al. 2014; Olalde, et al. 2014; Prufer, et al. 2014; Raghavan, et al. 2014; Skoglund, et al. 2014; Fu, et al. 2015; Gunther, et al. 2015; Cassidy, et al. 2016; Kilinc, et al. 2016; Omrak, et al. 2016; Schiffels, et al. 2016; Prufer, et al. 2017; Saag, et al. 2017; Gunther, et al. 2018). We counted the number of reads that mapped to regions around each of the three *AMY1* copies in the human reference genome (Usher, et al. 2015), and compared to bootstrap estimates of mean read depth with a linear correction for GC content (Methods, Figure 4, Supplementary Table 5). We find no significant difference between *AMY1* copy number in present day Europeans and ancient Europeans from the Iron Age/Medieval, Bronze Age, Neolithic, or Mesolithic periods. In particular, Mesolithic pre-agricultural hunter-gatherers have a mean copy number of 7.2 – not statistically different from present-day Europeans. Neolithic Early Farmers have a mean copy number of 8.4 – slightly higher, but not significantly different (at a Bonferroni-corrected significance level of 0.01) from present-day populations (P=0.02). We do find that four Upper Palaeolithic individuals dating between ~45,000-20,000 BP have lower *AMY1* copy number than present-day Europeans although, with a small sample size, this is also not statistically significant (mean 3.4, P=0.02). These results therefore suggest an expansion in copy number sometime earlier than ~10,000 BP and thus predating the development of agriculture.

**Figure 4:**
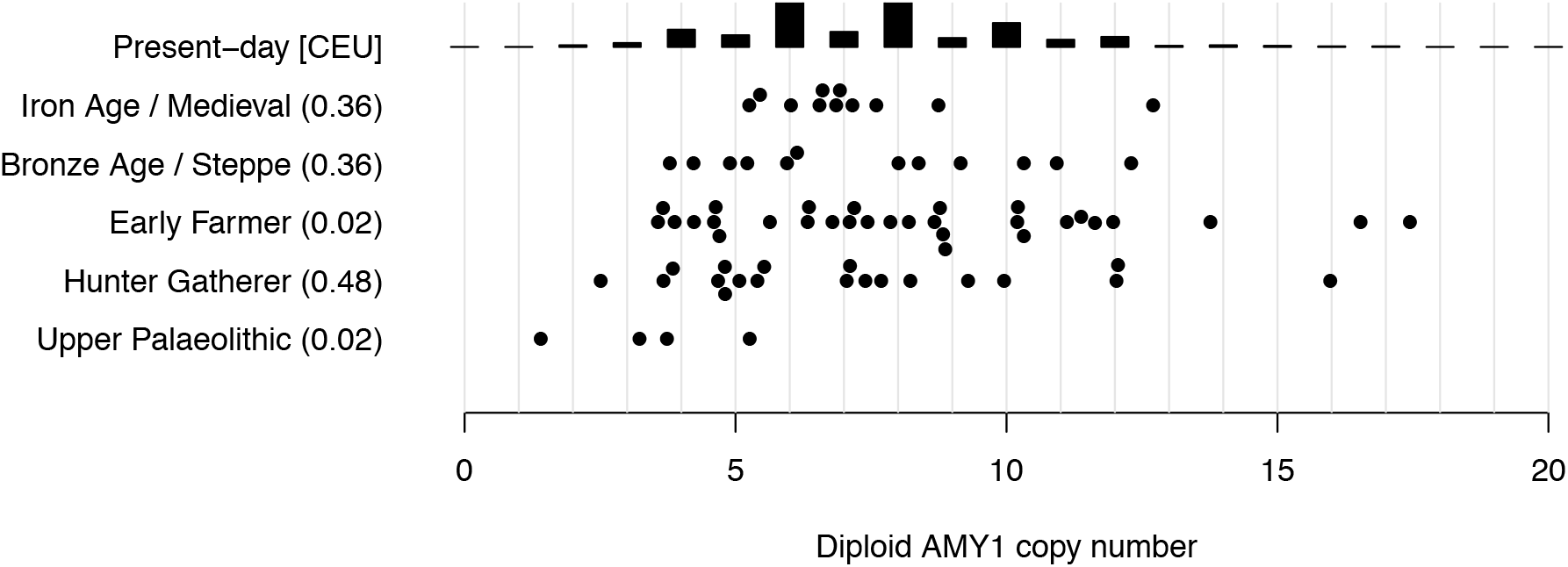
*AMY1* copy number. Inferred in ancient samples, arranged by time and subsistence strategy, compared with the distribution in a present-day population (CEU) with Northern European ancestry (Usher, et al. 2015). Parentheses: t-test P-values for difference between CEU and ancient populations.

Finally, we also investigated the history of other mutations that have been suggested to be involved in adaptation to agricultural subsistence. It has been proposed, based on both ancient and modern DNA, that the ergothioneine transporter gene *SLC22A4*–and in particular the nonsynonymous variant 503F (rs1050152)–was targeted by selection in Early Neolithic farming populations (Huff, et al. 2012; Mathieson, et al. 2015). However, analysis of our larger ancient DNA dataset reveals a more complicated story, with an allele frequency trajectory similar to that of the *FADS1* derived allele (Figure 5A). Specifically, the derived allele of rs1050152 is absent in hunter-gatherers, and present at low frequency in Early Farmers and Bronze Age populations. But, similar to the derived *FADS1* allele, it does not increase in frequency–and therefore does not appear to be under selection–in Early Farming populations. Strong selection on this allele likely only operated in the past few thousand years. This timescale is similar to the timescale over which the European lactase persistence variant (Enattah, et al. 2002) became common in Europe (Figure 5B) (Burger, et al. 2007; Allentoft, et al. 2015; Mathieson, et al. 2015). The “slow acetylator” variant of the *NAT2* gene has been hypothesized to be advantageous in agricultural populations (Luca, et al. 2008; Magalon, et al. 2008; Sabbagh, et al. 2011). However, we find that a SNP (rs1495741) that tags the “fast acetylator” phenotype (Garcia-Closas, et al. 2011) shows no change in frequency over the past 10,000 years (Figure 5C). Finally, we find that a variant (rs4751995) in the gene *PLRP2* that is relatively common in present-day populations with a cereal-based diet (Hancock, et al. 2010) is also more common in Early Farmers than hunter-gatherers, although it shows no evidence of selection in any of these populations (Figure 5D).

**Figure 5:**
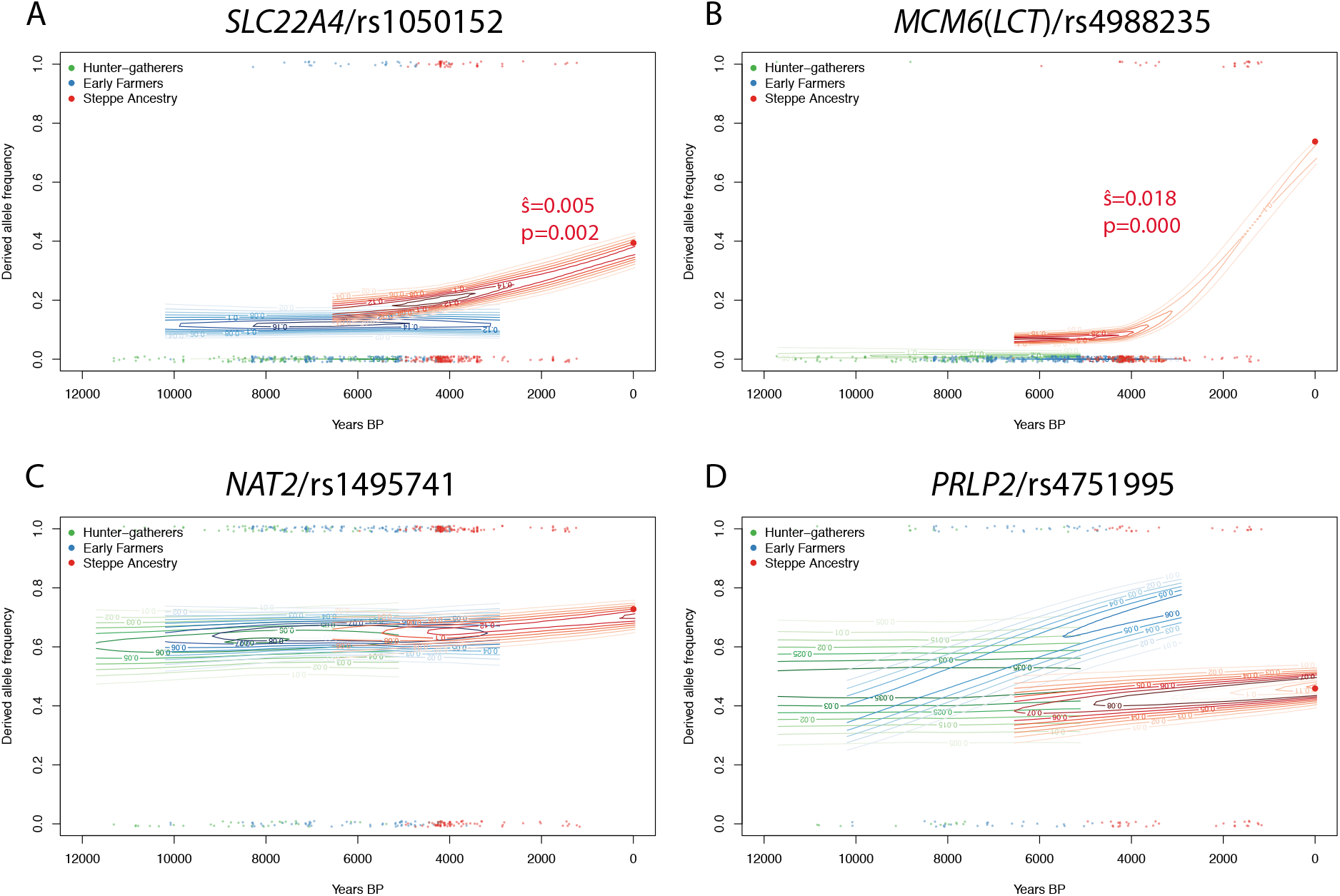
Allele frequency trajectories for other putative agricultural adaptation variants. As in Figure 2C, estimated allele frequency trajectories and selection coefficients in different ancient European populations. Significant selection coefficients are labelled.

## Discussion

The combination of high-quality genome sequence data from present-day people, and large ancient DNA datasets, provides new opportunities to investigate and understand the process of human adaptation. In particular, the temporal aspect of ancient DNA data adds another dimension to our analysis, allowing us to make precise inference of the timing of selection. We show that the derived allele was very rare or absent in all European populations, until it was introduced around 8500 BP by migration from early farmers carrying basal Eurasian ancestry. Although the derived allele is plausibly advantageous in populations that consume a plant-based diet, we find no evidence that it was actually selected in early European Farmers. Conversely, the low frequency of the derived allele in Upper Palaeolithic Europe is consistent with isotopic evidence that protein intake was dominated by animal protein at this time (Richards 2009). Finally, it is not known whether the Neanderthal allele would have had the same function as the derived modern human allele. But if it did, it would not support the claim that the derived allele is associated with a plant-based diet since Neanderthals, like Upper Palaeolithic modern humans, are thought to have consumed a largely meat-based diet (Bocherens 2009). Therefore, the derived/ancestral state at the major *FADS* haplotype may not be a simple marker of plant-vs. meat-based diet or more recent hunter-gatherer vs. farmer subsistence, but could reflect a more complex pattern of interaction based on unknown dietary and genetic factors.

Our results support and extend several recent results about the evolutionary history of the *FADS1* allele. It has previously been shown that the derived allele was anciently selected in Africa (Mathias, et al. 2012). We find that the ancestral allele was rare in early non-African populations, suggesting that there may have been selection for the ancestral allele as proposed by Ye, et al. (2017), but that it must have been before or during–rather than after–the out-of-Africa bottleneck. The fixation of the ancestral allele in present-day Native Americans has been interpreted as evidence for selection in their Siberian or Beringian ancestors (Amorim, et al. 2017; Harris, et al. 2017; Hlusko, et al. 2018). However, our analysis shows that this may not be the case, because the derived allele was likely very rare in Eurasia at the time of the Native American-Eurasian split 20-25,000 BP (Raghavan, et al. 2015). Both Ye, et al. (2017) and Buckley, et al. (2017) argue that recent selection in Europe for the derived allele was driven by changes in diet. Our results support this view, with the caveat that the relevant changes were not simply those associated with the Neolithic transition. Finally, Buckley, et al. (2017) found evidence for differential selective pressures across Europe and propose the marker rs174594 as the target of selection. ABC analysis is consistent with selection on rs174594 across Europe (Supplementary Table 6) but it is in strong LD with rs174546 and we were unable to test the model of Buckley, et al. (2017) further because rs174594 is not on the capture array that was used to generate most of the ancient DNA data we analyzed.

Similarly, due to limited ancient DNA data, we were not able to resolve the history of the *FADS1* allele in East Asia. ABC analysis was very sensitive to the exact demography that we assumed. When we capped recent *N_e_* at 45,000, we found that the ancestral allele was selected in the ancestors of present-day East Asians, although with a large credible interval (168,000-1,216,000 BP). The 39,000-year-old Tianyuan individual did not carry the derived haplotype, further suggesting that it was absent in the Upper Paleolithic ancestors of East Asians as well as Europeans. We estimated that the derived allele was selected more anciently than in Europe (i.e. ~35,300 BP with 95% CrI 9,400-108,000 BP). The most common East Asian derived haplotype is also an outgroup to the common European and African haplotypes (Supplementary Figure 2), which would be consistent with deriving from a separate, older, event. More ancient DNA from East Asia will help resolve this question, although we note that agriculture developed later in East Asia than in Western Eurasia, so it is likely that selection on the derived *FADS1* allele was also unassociated with the development of agriculture.

In the case of *FADS1* and all the other examples we investigated, the proposed agricultural adaption was either not temporally linked with the initial development of agriculture or showed no evidence of selection in Early Farmer populations. Instead, the three variants with any evidence of selection were strongly selected at some point between the Bronze Age and the present day, that is, in the past ~4000 years. This time period is one in which there is relatively limited ancient DNA data, and so we are unable to determine the timing of selection any more accurately. Future research should address the question of why this recent time period saw the most rapid changes in apparently diet-associated genes. One plausible hypothesis is that the change in environment or diet at this time was actually more dramatic than the earlier change associated with the initial development agriculture. For example, Early Neolithic populations may have retained some proportion of their diet from hunter-gatherer strategies, and only later transitioned to a completely agricultural diet. Other environmental factors like pathogen load or climate might also affect the selective pressures. Another hypothesis is that effective population sizes were so small, or populations so structured, before the Bronze Age that selection did not operate efficiently on variants with small selection coefficients. For example, analysis of present-day genomes from the United Kingdom suggests that effective population size increased by a factor of 100-1000 in the past 4500 years (Browning and Browning 2015). Larger ancient DNA datasets from the past 4,000 years will likely resolve this question.

## Materials and Methods

### Identifying and analyzing *FADS1* haplotypes

We defined derived haplotypes using the following procedure. Within the region ±50kb from rs174546 (hg19 chr11:61519830-61619830) there are 140 common (MAF > 0.05) SNPs when considering the SGDP (600 haplotypes) and archaic (6 haplotypes) samples, and restricting to sites where the ancestral allele can be determined based on the chimpanzee genome (PanTro2 Chimpanzee Sequencing Analysis Consortium (2005)). For each pair of SNPs within these 140, we compute the number of “mismatches” between ancestral and derived states. For example, if SNP 1 at haplotype *H* is in the ancestral state, but SNP 2 at *H* is in the derived state, then this counts as one mismatch. 1000 Genomes data was used to set the ancestral/derived state for each SNP (we used the chimpanzee allele if the allele was not present in 1000 Genomes). Counting up these mismatches for all pairs of SNPs, we obtain a 140 x 140 symmetric “mismatch” matrix *M*. We transform this matrix into an “adjacency” matrix *A* by setting each entry to 1 if the number of mismatches is below some threshold t, and 0 otherwise. In other words, if *M[i,j]* <= *t*, *A[i,j]* = 1, otherwise *A[i,j]* = 0. This adjacency matrix can then be interpreted as a graph, with SNPs as the nodes and edges between SNPs if they are connected (i.e. have a low number of mismatches).

From this graph, we find the largest clique (connected component where every pair of nodes is connected). This procedure can be interpreted as a way to find a subset of SNPs that are all in high LD with each other. The problem of finding the largest clique in a graph is NP-hard, but we use the Bron-Kerbosch algorithm which is more efficient in practice than brute force (Bron and Kerbosch 1973).

We use the procedure above to define 3 nested haplotypes: one considering all modern non-African populations (haplotype D), one considering all modern human populations (haplotype C) and one considering all modern and Neanderthal samples (haplotype B). For the mismatch thresholds, we use 12, 12, and 3 respectively (this last lower threshold accommodates the small sample size of archaic populations). Haplotype D contains 25 SNPs, haplotype C contains 11 SNPs, and haplotype B 4 SNPs. When there is more than one maximal clique of SNPs to choose from, we select one that is a subset of a larger core. This means that haplotype B is a subset of haplotype C and haplotype C is a subset of haplotype D. Note that our haplotype D differs slightly from the derived haplotype defined by Ameur, et al. (2012) which was 28 SNPs long.

We constructed a haplotype network for the haplotype B region from 1000 Genomes European, East Asian and African haplotypes, using the R package “pegas” (Paradis 2010). We inferred the phylogenetic relationship between the haplotypes by picking a single individual from the SGDP that was homozygous for each of the representative haplotypes and inferring the tree relating the haplotypes using BEAST2 (Bouckaert, et al. 2014). We rooted the tree with chimpanzee and used a uniform [6.6-10.0] million year prior for human-chimp divergence. This corresponds to a genome-wide mutation rate of ~4-6× 10^−10^ per-base per-year.

### ABC and startmrca analysis

To quantify the strength and timing of selection in different populations, we used approximate Bayesian computation (ABC), implemented in the *ABCtoolbox* package (Wegmann, et al. 2010). We use the pipeline implemented by Peter, et al. (2012), which uses a model selection approach to distinguish selection on a *de novo* mutation (SDN) from selection on standing variation (SSV). Intuitively, this approach simulates data under both the SDN and SSV models, for a range of parameters, and then selects the simulations that best match the observed data, in the sense of being close in the space of a set of predefined summary statistics. The parameters of the selected simulations are then used to estimate a posterior distribution for the parameters of interest. In addition to SDN/SSV model selection, we estimated two continuous parameters: the selection-onset time and the selection coefficient. To perform the simulations for ABC we used *mbs* (Teshima and Innan 2009), which creates a selected allele frequency trajectory backward in time from a specified present-day frequency. This implicitly creates a range of selection-onset times in the past, which we used as a prior. For the selection strength, we used a uniform prior of [-4,-1] on the base-10 log of the selection coefficient.

We chose the derived allele (C) of rs174546 as the putatively selected allele, and analyzed 50kb on either side for a total region length of *L*=100kb. We fixed the mutation rate at 1.25×10^−8^ per base per generation, and used the recombination rate from the combined HapMap 2 map (International HapMap Consortium 2007). Since the recombination map may have changed over time, we also repeated the analysis using a constant recombination rate of 1.1785×10^−8^ per base per generation (the average rate in this region). We also set the current effective population size *N_e_* to 10,000, and the heterozygote advantage at 0.5. This mutation rate is at the low end of human mutation rate estimates (Scally 2016), but a higher mutation rate would only lead to more recent estimates for the onset of selection. For each region (AFR, EAS, and EUR) and each model (SDN and SSV), we simulated 1 million datasets. Each dataset had a sample size of *n*=170 haplotypes and a length of *L*=100kb to match the 1000 Genomes data. Within each region we chose five representative populations for further analysis: ESN, GWD, LWK, MSL, and YRI for AFR; CDX, CHB, CHS, JPT, and KHV for EAS; and CEU, FIN, GBR, IBS, and TSI for EUR. Based on the current frequency of the selected allele in the subpopulations for each region, we computed simulation priors: [0.984, 0.992] for AFR, [0.304, 0.670] for EAS, and [0.575, 0.699] for EUR. We also used population-specific demographies from PSMC, based on Yoruba (AFR), Han (EAS), and French (EUR) individuals from the SGDP (Mallick, et al. 2016).

For each simulated dataset we computed 27 summary statistics (as described in Peter, et al. (2012)). During the ABC estimation phase, we retained the top 500 simulated datasets with statistics closest to each real dataset (15 in total, one for each subpopulation), and computed posterior distributions for the selection-onset time and selection coefficient. We computed combined estimates (Supplementary Table 2 and 3) by assuming a lognormal distribution for the posteriors and averaging over the five population-specific estimates. Finally, we averaged over the two (SDN/SSV) model tested.

For EAS and EUR, we also wanted to test for ancient selection on the ancestral allele (T) of rs174546. To this end, we merged the subpopulations for both EAS and EUR, and selected 295 haplotypes with the ancestral allele for each. Then we ran our ABC procedure in the same way as before, except with a “current” allele frequency of 0.999999 (*mbs* does not allow 1 for technical reasons). The EAS results were very sensitive to demography–particularly to the value of present-day *N_e_*–so we restricted maximum *N_e_* to the value of 45,000 used by Gravel, et al. (2011).

The main limitations of the ABC approach are that it does not model complex modes of selection and relies on simulating under an appropriate range of parameters. We checked that ancient selection would not obscure more recent signals of selection by simulating data under a realistic model of alternating selection coefficients using *SLiM2* (Haller and Messer 2017), and rerunning the ABC analysis (Supplementary Figure 5). Similarly, the results are conditional on simulating under the correct demography and with an appropriate range of parameters. For our key results, we checked that the distributions of simulated summary statistics were broadly consistent with the observed values, indicating that our simulations are reasonable (Supplementary Figures 6-8). When we modeled selection on the ancestral allele, we noticed that the observed values of the Fay and Wu H statistics were often outside the simulated distribution (Supplementary Figure 7) – most likely because we flipped the selected allele – so we excluded these four statistics from the ABC estimation for selection on the ancestral allele.

*Startmrca* (Smith, et al. 2017) is a method for estimating the selection-onset time of a beneficial allele. It uses an HMM (in the “copying” style of Li and Stephens (2003)) to model present-day haplotypes as imperfect mosaics of the selected haplotype and a reference panel of non-selected haplotypes. We used this method to estimate the selection-onset time for the ancestral allele of rs174546 in the EAS and EUR superpopulations described above. We used YRI individuals that are homozygous for the derived allele as the reference panel. Following Smith, et al. (2017), we used a 1Mb region surrounding rs174546, the HapMap combined recombination map in this region (International HapMap Consortium 2007), 100 haplotypes in the selected population, and 20 haplotypes in the reference population. To obtain selection-onset time estimates for each population, we ran 5 independent MCMC chains, each with 10,000 iterations. We post-processed the results by discarding the first 6000 iterations (burn-in), and retaining the remaining successful iterations over all 5 chains. To obtain credible intervals, we took the 2.5 and 97.5 quantiles of each resulting distribution (Supplementary Table 3.2).

We also analyzed selection for the derived allele of rs174546 in the EAS and EUR superpopulations. In this case, there were not enough AFR individuals homozygous for the ancestral allele to use as a reference population. So instead we used a “local” reference population, i.e. EAS and EUR individuals homozygous for the ancestral allele. We also use a lower prior for the derived allele ([0-4,000] generations vs. [0-20,000] for the ancestral). We did not analyze AFR because the selection is likely too old for the method, and there is no suitable reference population. We note that *startmrca* tends to underestimate the time of very ancient selective events (the “star” genealogy assumption becomes less appropriate). It also does not account for selection on standing variation, which would result in an overestimation of the selection-onset time, so our *startmrca* quantitative results are likely unreliable.

### Ancient DNA analysis

We first analyzed 52 Palaeolithic and Mesolithic samples (Fu, et al. 2014; Jones, et al. 2015; Fu, et al. 2016; Sikora, et al. 2017) for the presence of the derived *FADS1* allele. These samples were typed on a capture array (“1240k capture”) that contains 5 of the 25 SNPs that define haplotype D. We divided the samples into population groups as defined by Fu, et al. (2016) and inferred the allele frequency in each of these populations by maximizing the following likelihood function:

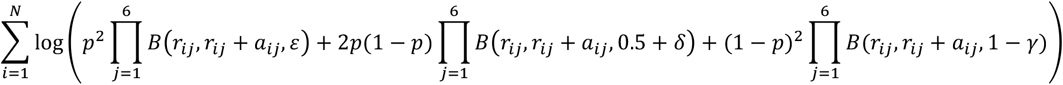

where *r_ij_* and *a_ij_* are the number of reference and alternative reads from individual *i* at SNP *j, N* is the number of individuals in the population. *B(x,n,p)* is the binomial probability of seeing *x* successes out of *n* trials with probability *p*, and *ε*,δ,γ are small error probabilities, which we set to 0.1, 0, 0.1 for transversions, 0.15, 0.05, 0 for C>T or G>A transitions, and 0, 0.05, 0.15 for T>C or A>G transitions. This implies a conservative 10% rate of contamination or error, and a 5% deamination rate. We computed binomial confidence intervals assuming an effective sample size of 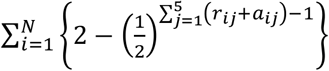.

To impute the *FADS1* haplotype in the Upper Palaeolithic and Mesolithic samples, we computed genotype likelihoods at each typedSNP in the 5Mb region around rs174546, for each individual, assuming a binomial distribution of reference and alternative allele counts and a 5% deamination rate. We then imputed using *BEAGLE 4.1* (Browning and Browning 2016), and the 1000 Genomes reference panel downloaded from bochet.gcc.biostat.washington.edu/beagle/1000_Genomes_phase3_v5a. We filtered out imputed SNPs with a genotype probability of less than 0.8 and used the remaining SNPs to determine the presence of the allele. In order to estimate the false positive rate, we simulated read data at different coverages for 50 individuals from the 1000 Genomes EAS (East Asian) super-population who are homozygous for the ancestral allele. We computed genotype likelihoods and imputed as for the ancient data, having first removed the 50 test individuals from the reference panel. We performed 10 simulations for each coverage level and used the frequency at which the derived allele was imputed as an estimate of the false positive rate.

To analyze the Holocene history of *FADS1* and other alleles, we assembled a dataset of 1078 published ancient samples, most of which were typed on the “1240k” capture array which targets ~1.24 million SNPs. We used the pseudo-haploid version of these data, where each individual has a single allele at each SNP, from a randomly selected read. We classified these individuals into “hunter-gatherer”, “Early Farmer” and “Steppe ancestry” populations as follows. First, we ran supervised ADMIXTURE (Alexander, et al. 2009) with K=4, and the four populations: WHG, EHG, Anatolia_Neolithic and Yamnaya_Samara fixed to have cluster membership 1, as previously described (Mathieson, et al. 2018). We then classified individuals based on their inferred ancestry. If they had more than 25% ancestry from the Yamnaya_Samara cluster and dated later than 6000 BP, we classified them as “Steppe ancestry”. If they had less than 25% ancestry from the Yamnaya_Samara cluster, more then 50% from the Anatolia_Neolithic cluster and dated earlier than 2,000 BP, we classified them as “Early Farmer”. Finally, if they had less than 25% ancestry from the Yamnaya_Samara cluster and less than 50% ancestry from the Anatolia_Neolithic cluster and were dated earlier than 5100 BP, we classified them as “hunter-gatherer”. We excluded 23 samples that did not fit into any of these classifications, leaving an analysis dataset of 1055 samples of which 669 had coverage at rs174546. These classifications are informed by previous analysis that combined genetic, chronological and archaeological information, and largely correspond to classifications that would be derived from archaeological context alone. We estimated allele frequency trajectories and selection coefficients separately for each population using a method that fits a hidden Markov model to the observed frequencies (Mathieson and McVean 2013), assuming an effective population size (*N_e_*) of 10,000 in each population. We also analyzed separately the ancient individuals from Northern and Southern Europe (Supplementary Figure 3). For this analysis, we defined “Northern Europe” to be the region east of 13°E and north of 45°N, or west of 13°E and north of 49°N, with “Southern Europe” as the complement. This corresponds approximately to the region reached by the Neolithic transition by 7,000 BP (Fort 2015).

To call *AMY1* copy number we assembled a set of 76 ancient genomes with shotgun sequence data that had nonzero mapped coverage at the locus. The majority of published ancient shotgun genomes have zero coverage, presumably because the copy number variable region was masked during alignment. We counted the number of reads that mapped to the any of the three *AMY1* duplicate regions in the human reference genome (Usher, et al. 2015) and compared the total to the average read depth in 1000 random regions of chromosome 1, of the same size as the *AMY1* duplicate regions. We fitted a linear model of coverage as a function of GC content to these 1000 regions, for each individual, and used this to correct our estimates for GC bias.

### Analysis of lipid GWAS hits

We tested the directionality of lipid-associated alleles using genome-wide association meta-analysis results for LDL, HDL and TG (Teslovich, et al. 2010). Specifically, we constructed a list of SNPs with P-values below a given cutoff by iteratively selecting the SNP with the lowest P-value and then removing all SNPs within 500kb. For each of these SNPs we extracted allele frequencies in the EUR and AFR super-populations and then tested whether trait increasing alleles were more common in AFR than EUR (Figure 3A).

To analyze the LDL hits in ancient samples, we first identified all SNPs with an association P-value less than 10^−6^ that were on the capture array used to genotype the majority of the ancient samples. We iteratively removed SNPs within 250kb of the most-associated SNPs to produce an independent set of associated SNPs. For each (pseudo-haploid) individual, we constructed the “LDL score” by counting the proportion of these SNPs at which that individual carried the trait-increasing allele (Figure 3B). We found no significant differences with respect to ancestry when we fitted a binomial generalized linear model with ancestry as a covariate. We also fitted a model including date as a covariate to test for significant differences over time, also with a nonsignificant result.

## Acknowledgments

We thank Rasmus Nielsen, Joshua Schraiber and two anonymous reviewers for helpful comments on an earlier draft, and Benjamin Peter for help running the ABC methods.

